# Auditory experience prevents loss of the innate song preference as a selective cue in *Drosophila*

**DOI:** 10.1101/2020.07.16.205443

**Authors:** Xiaodong Li, James Kruszelnicki, Yukina Chiba, Hiroshi Ishimoto, Yuki Ishikawa, Azusa Kamikouchi

## Abstract

Auditory learning is a prerequisite step for acoustic communication learning, which was previously assumed to be restricted to animals with high levels of cognition, such as humans, cetaceans, and birds. How animals that rely on auditory learning for acoustic communication form sound preferences is not known. Fruit flies are a recently proposed novel animal model for studying experience-dependent auditory perceptual plasticity because of their ability to acquire song preferences via song exposure. Whether fruit flies have innate courtship song preferences, however, is unclear. Here we report that, similar to songbirds, fruit flies exhibit an innate preference for conspecific courtship songs. Maintenance of innate song preference requires song input, reminiscent of the song learning process in songbirds. Our findings also indicate that the response to conspecific and heterospecific songs manifests temporal and experience-dependent differentiation, which may underlie innate song preference and its plasticity. In addition, we find that flies have a robust ability to reacquire song preference during aging. Fruit flies thus offer a novel and simple approach for studying sound preference formation and its underlying mechanisms.

## Introduction

Auditory perceptual preference is prerequisite for social acoustic communication. Auditory learning does not originate from a *tabula rasa* in either humans or songbirds. Infant humans are capable of discriminating all phonetic contrasts of existing languages (**1**), and already show categorical perception of sounds before learning (**2, 3, 4**). Young songbirds innately prefer to respond to and learn conspecific songs over heterospecific songs (**5, 6, 7, 8, 9, 10**). These findings clearly demonstrate an innate predisposition of auditory perception. On the other hand, numerous studies suggest that these innate predispositions can be modified by experience. Human infants prefer the voices of their mothers over others, speech over non-speech sounds, and native over non-native languages (**11, 12, 13**). By the end of their first year, infants retain distinctions of the phonetic contrasts in their native language, while losing such distinctions in non-native languages (**4, 14**). The interaction between innate predisposition and auditory learning in establishing auditory perceptual preference, however, remains unclear.

Three different models are used for studying perceptual learning in humans and birds to examine how sensory experience alters initial predisposition: selective, instructive, or a mixture of both (**4, 15**). A purely selective model suggests that extensive innate knowledge exists in the brain to guide future learning. The core of this model is that in forming memory, the neural circuitry exhibits highly selective responses to conspecific sounds and experience is required to activate this circuitry. In a purely instructive model, there is no predisposition and animals are open to learn any sound to which they are exposed. The third model combines instruction in the memory phase and selection in the production phase. Knowing whether auditory experience supplies a selective or instructive cue in this process is critical to our understanding of how innate predispositions and experience interact in shaping perceptual auditory preference.

Song learning processes in songbirds share many similarities with language acquisition in humans. To acquire a song, the process by which young songbirds hear and memorize the auditory cues of adults (auditory learning) necessarily precedes the process of producing learned sounds (sensorimotor learning) (**4**). In songbirds of both sexes, song perception is shaped by early auditory experience. On the basis of exposure to a tutor’s song, males learn to sing and females learn to evaluate male songs during courtship (**16, 17, 18**). For males, it is clear that song memory serves as a template for matching in vocal learning, while for females the significance of song memory is unclear. One possibility is that song memory guides female attraction to songs with specific features during mate choice by sexual imprinting (**19, 20, 21, 22**).

A report consistent with this notion comes from an auditory learning study in the fruit fly *D. melanogaster*, in which the song preference of flies is shaped by an experience of hearing particular conspecific songs (**23, 24**), in contrast to the traditional view that all steps in mating (including sensory perception) are innate (**25, 26, 27**). As the majority of *D. melanogaster* males are able to successfully mate within 2 days after eclosion (**28**), and immature males are also mating targets for mature males (**29, 30**), it is possible that most males hear courtship songs from peers and this experience shapes their auditory preference (song preference learning). It is noteworthy that the song preference of flies is shaped only by their exposure to conspecific songs, and not to heterospecific songs (**24**). Therefore, although Li et al. (**24**) proposed the idea that auditory perception in flies is not genetically hard-wired but rather experience-dependent, it remains to be investigated whether flies have an innate predisposition for song preference and, if so, how the interaction between the innate predisposition and auditory experience shapes song preference.

Here, we used behavioral manipulations to show that *D. melanogaster* has an innate song preference, and that conspecific song experience serves as a selective cue, rather than an instructive cue, to prevent loss of the innate song preference. We further show that flies have a long sensitive period during which they are able to repeatedly reacquire auditory preference. The results of this study, together with previous research (**24**), are consistent with the selective model for auditory learning in at least 3 aspects. First, flies innately exhibit larger auditory responses to conspecific songs than to heterospecific songs. Second, flies cannot sustain song preference during song deprivation or social isolation. Third, flies selectively learn from song experience to prevent loss of the innate preference. These features pave the way for using fruit flies as a model to investigate the conserved neural mechanisms underlying auditory preference and learning in insects and vertebrates.

## Results

### Young adult *Drosophila* show innate song preference

The biased song preference learning of 7-day-old flies, in which their preference is shaped by exposure to conspecific song but not to heterospecific song (**24**), led us to investigate whether flies have an innate song preference. We asked if younger male flies exhibit innate song preference irrespective of song experience. When we housed the young males in a chamber, we detected spontaneous chasing behavior and unilateral wing extension between males (**Movies S1-S3** for 2, 3, and 4 day-old males; **Fig. 1*A***, right side), which are typical steps in the courtship ritual of fruit-fly males (**31**). This observation confirmed that group-reared intact males are exposed to conspecific courtship song during homosexual courtship and thus they get auditory experience in the absence of females. In the following experiments, therefore, wing-intact flies housed in a single-sex group are referred to as experienced flies. On the other hand, flies whose wings were clipped soon after eclosion are referred to as naive flies, because wing-clipped males cannot emit wing-beat sounds. We defined 4-day-old flies as young flies and tested their song preference after conditioning in 1 of 3 groups (**Fig. 1*A***). To test the song preference, we used the chaining assay to assess the chasing behavior of male flies evoked by a courtship pulse song (**23, 32**). Two types of artificial pulse songs, one with a 35-ms inter-pulse interval (IPI) and the other with a 75-ms IPI, were used to represent conspecific and heterospecific songs, respectively (**24**).

**Fig. 1.**
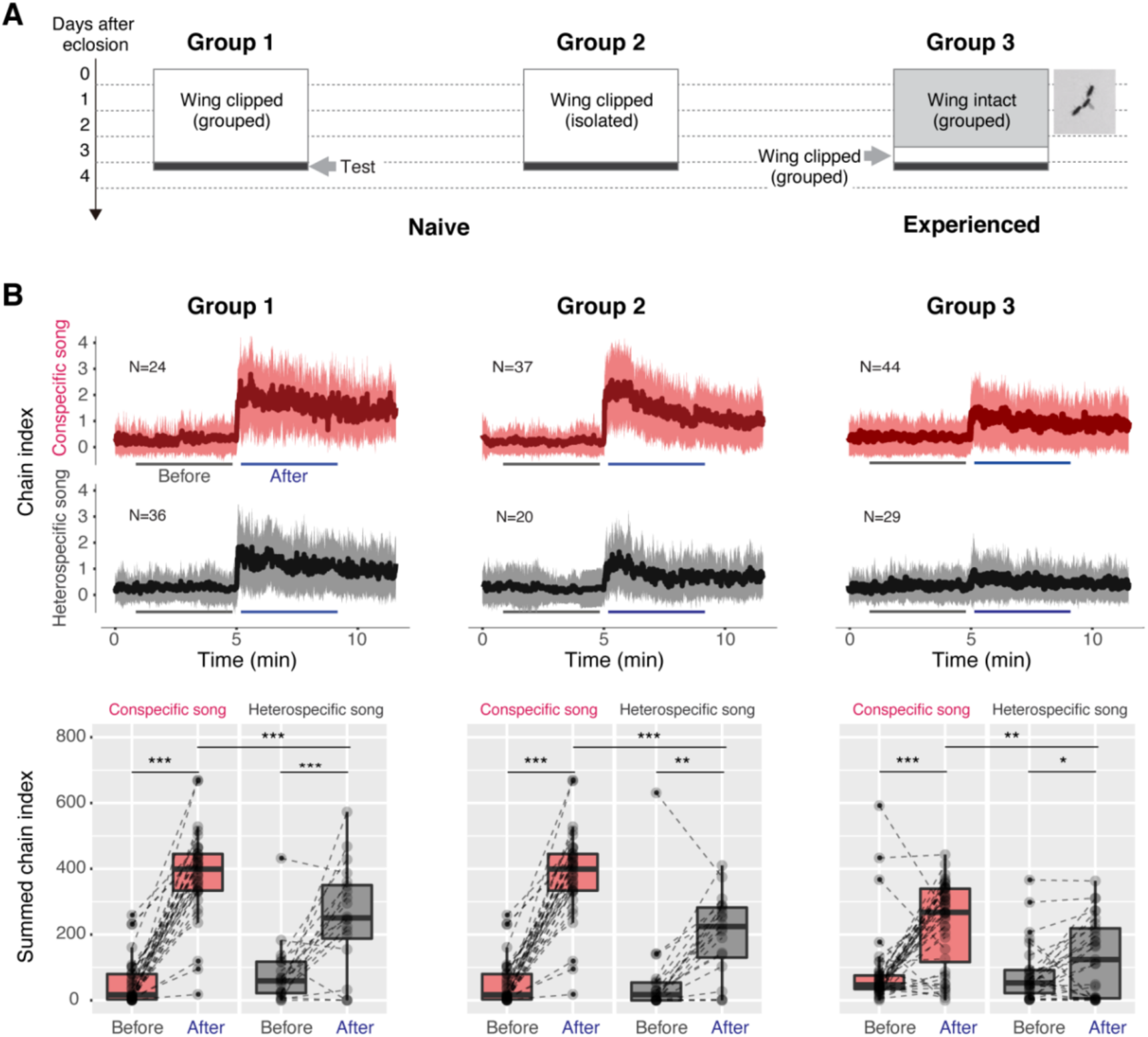
Young adult flies exhibit innate song preference. (*A*) Flies were housed for 3 days under different experimental conditions; grouped without wings (Group 1), isolated without wings (Group 2), and grouped with intact wings (Group 3). The image on the right side shows the homosexual courtship between males. Each horizontal dashed line indicates ZT = 0. Refer to **Table S1** for the experimental conditions. (*B*) Time-courses of the chain index in response to playback of the conspecific song (red) and heterospecific song (grey). Sound playback starts at 5 min. The bold line and ribbon represent the mean value and standard error, respectively. The box plot shows the summed chain index in a 4-min window before or after the song start (indicated by the horizontal lines). N.S., not significant, *P*>0.05; \**P*<0.05; \*\**P*<0.01; \*\*\**P*<0.001 Mann-Whitney U tests with the Benjamini–Hochberg correction and Wilcoxon signed-rank test with the Benjamini–Hochberg correction were used in multiple comparisons of mixed samples. N, number of behavioral chambers examined. Refer to **Table S2** for details of statistical tests and scores.

By performing chaining assays, we found that young naive flies reared in grouped conditions (Group 1, **Fig. 1*A***) preferred to respond to the conspecific song over the heterospecific song (**Fig. 1*B*** left panel). To exclude other sensory cues from peers in grouped conditions, we tested singly-reared wing-clipped males, also regarded as naïve flies (Group 2, **Fig. 1*A***), and again observed a clear preference for the conspecific song (**Fig. 1*B*** middle panel). Experienced male flies reared with intact wings in grouped conditions (Group 3, **Fig. 1*A***) exhibited the same preference (**Fig. 1*B*** right panel). The consistent biases in these 3 conditions indicate that social environment or hearing experience does not affect the song preference of young adult males, suggesting a genetically innate preference of male flies toward conspecific songs.

### Recovery of the heterospecific song response induces loss of song preference

In previous studies, we reported that 6 to 7 day-old naive flies did not show such song preference, although they could acquire this preference after auditory exposure (namely, song preference learning) (**23, 24**). We defined 6 to 7 day-old flies as mature flies. Further comparison between young naive flies (Group 1 in this study) and mature naive flies (data shared by Li et al. [**24**]) revealed that the response to the conspecific song was equivalent (**Fig. S1*A***), while the response to the heterospecific song was weaker in young naive flies (**Fig. S1*B***). These results indicate that the auditory response to songs with these two IPIs manifests temporal differentiation, and the change in the song preference in different time windows is induced mainly by the varying response to heterospecific song.

In these experiments, naive male flies were deprived of courtship song sources and thus these results raised the hypothesis that prolonged song deprivation results in the loss of innate song preference in male flies. We evaluated this hypothesis by examining whether song preference in flies maintained at the mature adult stage is lost if flies were deprived of song input. To achieve this, we housed intact flies in a grouped condition for 5 days after eclosion and then clipped their wings. These wing-clipped males were reared in grouped conditions for either 2 days, 3 days, or 4 days before being tested in chaining assays (Groups 4, 5, & 6, respectively, **Fig. 2*A***). During the latter period, the males should have been deprived of any song input. We found that song preference resulting from a 5-day grouped experience (**24**) was sustained for up to 3 days (**Fig. 2*B*** left & middle panels), but disappeared after 4 days of song deprivation (**Fig. 2*B*** right panel). These findings indicate that song preference maintained during the mature adult stage can be lost via song deprivation.

**Fig. 2.**
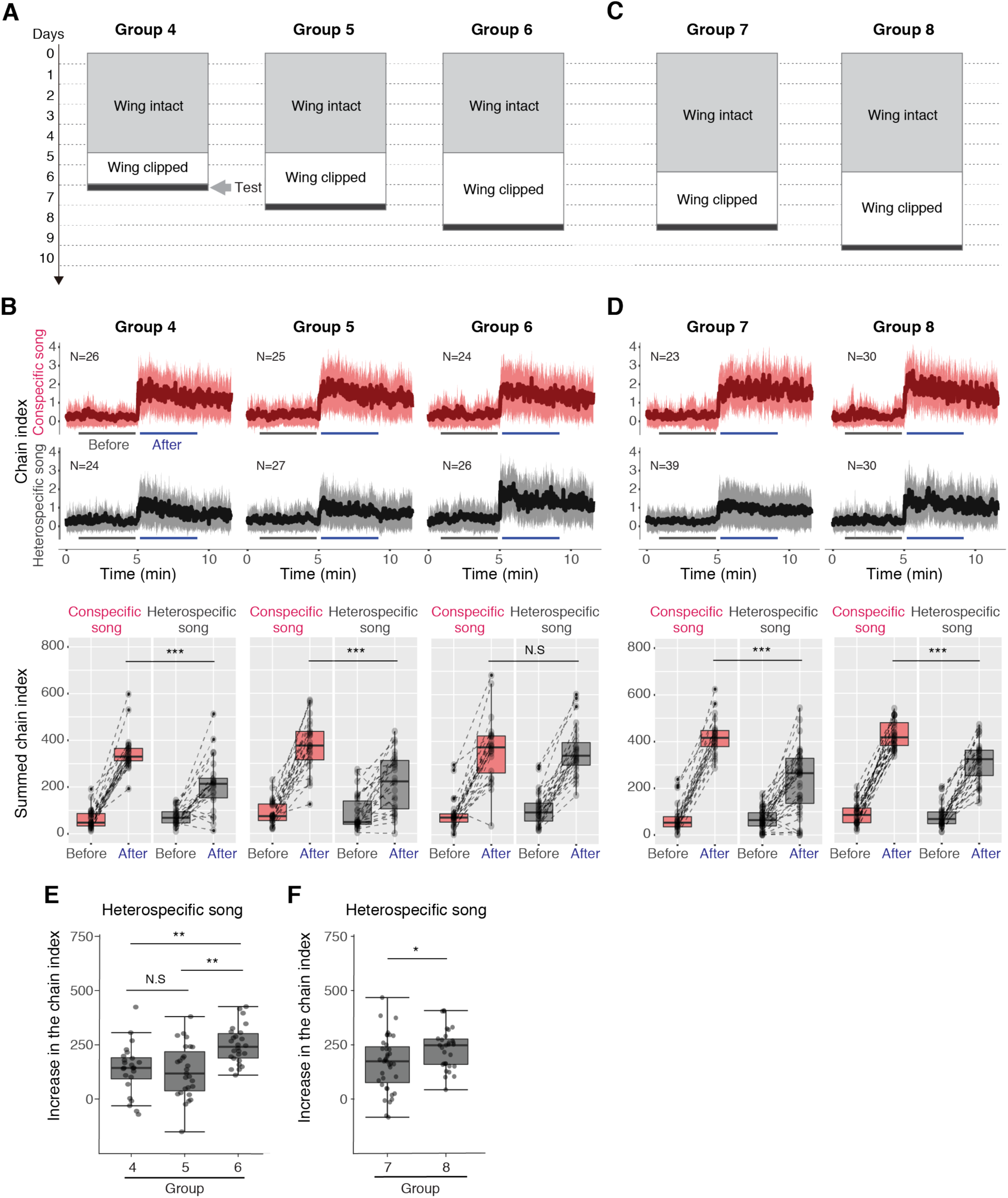
Song preference is lost after song deprivation. (*A*) Wing-intact male flies were reared in grouped conditions for 5 days and wing-clipped conditions for 2 days (Group 4), 3 days (Group 5), or 4 days (Group 6) before the chaining assays. Refer to **Table S1** for the experimental conditions. (*B, D*) Time-courses of the chain index in response to playback of the conspecific song (red) and heterospecific song (grey). (*C*) Wing-intact male flies were reared in grouped conditions for 6 days and wing-clipped conditions for 3 days (Group 7) or 4 days (Group 8) before the chaining assays. (*E, F*) Increase in the chaining activities induced by the heterospecific song for each group. Increase in the chain index following song onset = (summed chain index in 311-550 s) – (summed chain index in 51-290 s). N.S., not significant, *P*>0.05; \**P*<0.05; \*\**P*<0.01; \*\*\**P*<0.001 Mann-Whitney U tests with the Benjamini–Hochberg correction and Wilcoxon signed-rank test with the Benjamini–Hochberg correction were used in multiple comparisons of mixed samples; Kruskal–Wallis tests followed by the Mann-Whitney U test with the Benjamini–Hochberg correction was used in multiple comparisons of independent samples; Mann-Whitney U tests were used for 2-sample comparisons. N, number of behavioral chambers examined. Refer to **Table S2** for details of statistical tests and scores.

We then investigated whether a longer grouped condition could prolong the maintenance of song preference. We kept intact males in grouped conditions for 6 days rather than 5 days, and then clipped their wings and deprived them of song input for either 3 or 4 days before testing them in chaining assays (Groups 7 and 8, **Fig. 2*C***). Song preference maintained from the 6-day grouped condition remained even after 4 days of song deprivation (**Fig. 2*D***). This suggests that longer song experiences within grouped conditions prolong maintenance of the song preference.

We asked whether the disappearance of song preference after 4 days of song deprivation in the Group-6 flies was induced by an increase in the heterospecific song response or a decrease in the conspecific song response. We quantified the increase in the chain index following song onset in the chaining assays (see **Materials and methods**). Interestingly, for flies with 5 days of group experience (**Fig. 2*A***) the increase in the chain index in response to the heterospecific song did not differ between Groups 4 and 5, i.e., 2 or 3 days after song deprivation (**Fig. 2*E***), but significantly increased after 4 days of song deprivation (Group 6 in **Fig. 2*E***). For flies with 6 days of group experience (**Fig. 2*C***), the increase in the chain index in response to the heterospecific song also significantly increased from 3 days of song deprivation to 4 days (**Fig. 2*F***). On the other hand, we found that the increase in the chain index in response to the conspecific song did not change across groups (**Fig. S2 *A*** and ***B***). These findings indicate that the response to heterospecific song recovers following song deprivation, while the response to conspecific song remains invariable, thus leading to a disappearance of song preference after 4 days of song deprivation for flies with 5 days of group experience (Group 6). It is noteworthy that for flies with 6 days of group experience, although the response to the heterospecific song tended to increase from 3 days of song deprivation to 4 days (**Fig. *2F***), this recovery was not sufficient to abolish the song preference seen in Group 8 flies (**Fig. *2D***). Thus, these findings suggest that longer exposure to conspecific song enables longer retention of song preference.

The above results demonstrate that varying song preference can be induced by temporal differentiation of the auditory response to conspecific and heterospecific songs, and that the song preference can be lost due to song deprivation. Previous research showed that although mature flies kept in wing-clipped conditions lost the song preference, those with intact wings in grouped conditions or wing-clipped but exposed to artificial conspecific songs after eclosion maintained their song preference (**24**). These results, together with our finding shown in **Fig. 2*E***, indicate that conspecific song input suppresses the auditory response to heterospecific songs, and removing the conspecific song input results in the recovery of the heterospecific song response, suggesting that conspecific song input is required to maintain innate song preference. Together, these results support the theory that auditory experience reinforces genetic innate preference.

### Hearing experience in the first 3 days after eclosion suppresses responses to conspecific songs

Previous research revealed that the conspecific song response in mature male flies is always sustained, even after long-term conspecific song exposure (**24**). We noticed, however, that auditory experience affects the response of young flies to both conspecific and heterospecific songs; compared with the young naive flies (Group 1), young flies with intact wings for 3 days (Group 3) had a reduced response to both songs (**Figs. 1** and **S3**).

As young male flies are courted by other males after eclosion (**29, 30**; **Movies S1- S3**), we hypothesized that the song experience in young males suppressed the song response. To test this, we utilized the sound training setup **(32**) to expose wing-clipped male flies to artificial sounds in isolated conditions for the first 3 days after eclosion. We used white noise, pure tone, or conspecific song as the training sounds and then performed chaining tests at 4 days after eclosion (Groups 9, 10 & 11, respectively, in **Fig. 3*A***). Although the flies in all of the groups exhibited a strong response to the conspecific song (**Fig. 3*B***), only Group 11 flies that were trained with the conspecific song had reduced responses compared with the young naive isolated flies (Group 2; **Fig. 3*C***). This markedly reduced response was not detected in Group 9 and 10 flies, which were exposed to white noise or pure tone (**Fig. 3*C***). A previous report that tested the response in mature flies, however, found that 6-day exposure to the conspecific song did not suppress their auditory response to conspecific songs (**24**). These two findings together suggest that the response to conspecific song becomes more resistant to change during maturation. Moreover, we found that the response to the conspecific song was not suppressed in the group with a delayed 3-day exposure to the conspecific song (Group 12, **Fig. 3*C***), nor in the group with 3 days of exposure followed by 3 days of recovery (Group 13, **Fig. 3*C***).

**Fig. 3.**
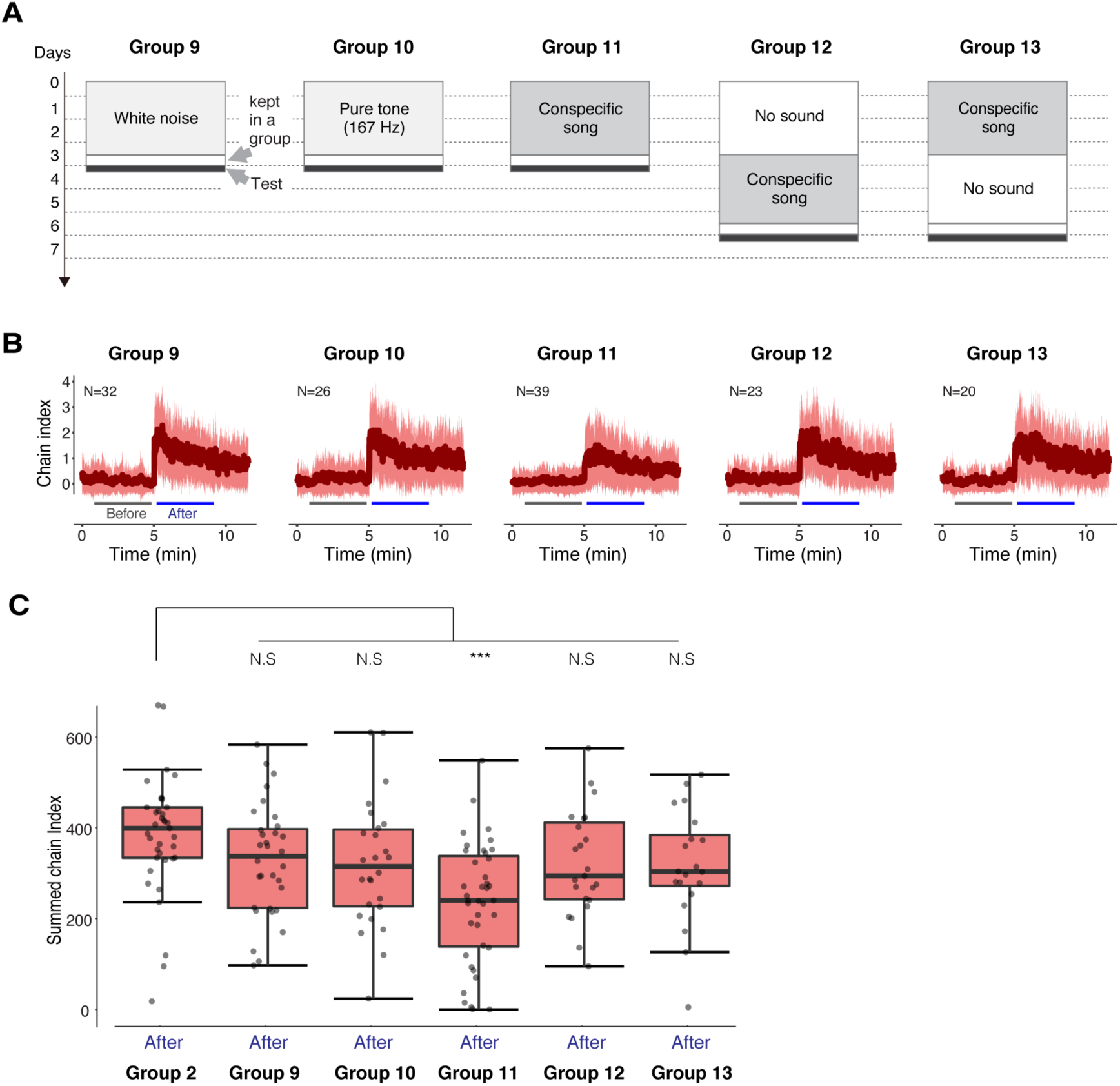
Response to conspecific song is plastic in young adult flies. (*A*) Wing-clipped flies were exposed to white noise (Group 9), pure tone (Group 10), or conspecific song (Group 11) for the first 3 days after eclosion prior to the chaining tests. The onset of the conspecific song exposure for wing-clipped flies in Group 12 was delayed for 3 days. Flies in Group 13 were allowed to recover for 3 days after a 3-day exposure to the conspecific song. Refer to **Table S1** for the training conditions. (*B*) Time-courses of the chain index in response to playback of the conspecific song. (*C*) Summed chain index in the 4-min window after song onset in Groups 9 to 13 were compared with that in Group 2 in response to the conspecific song. N.S., not significant, *P*>0.05; \**P*<0.05; \*\**P*<0.01; \*\*\**P*<0.001 Wilcoxon signed-rank test were used to compare two dependent samples; Kruskal–Wallis tests followed by the Mann-Whitney U test with the Benjamini–Hochberg correction was used in multiple comparisons of independent samples. N, number of behavioral chambers examined. Refer to **Table S2** for details of statistical tests and scores.

Taken together, the above results indicate that exposure to the courtship song within the first 3 days of eclosion suppresses auditory responses to the conspecific song, suggesting that the auditory response to conspecific song is also plastic when flies are young.

### Auditory exposure restores lost song preference

The possibility of restoring phonetic discrimination ability in individuals who have lost this capability is of great interest (**34, 35, 36**); we therefore asked if it is possible to restore lost song preference in *D. melanogaster*. Flies housed alone for 5 days after eclosion do not show song preference (**24**). We thus tested whether the song preference of the flies was restored if after 5 days of isolation they were housed in group conditions. To test this, we established 4 fly groups (**Fig. 4*A*** and **Table S1**), each of which was reared in different combinations of singly and group-housed conditions. The first 3 groups, Groups 14, 15, and 16, represent control groups and the other, Group 17, was the experimental group. In all of the groups, intact flies were kept for 10 days and their wings were clipped 1 day before the chaining assay.

**Fig. 4.**
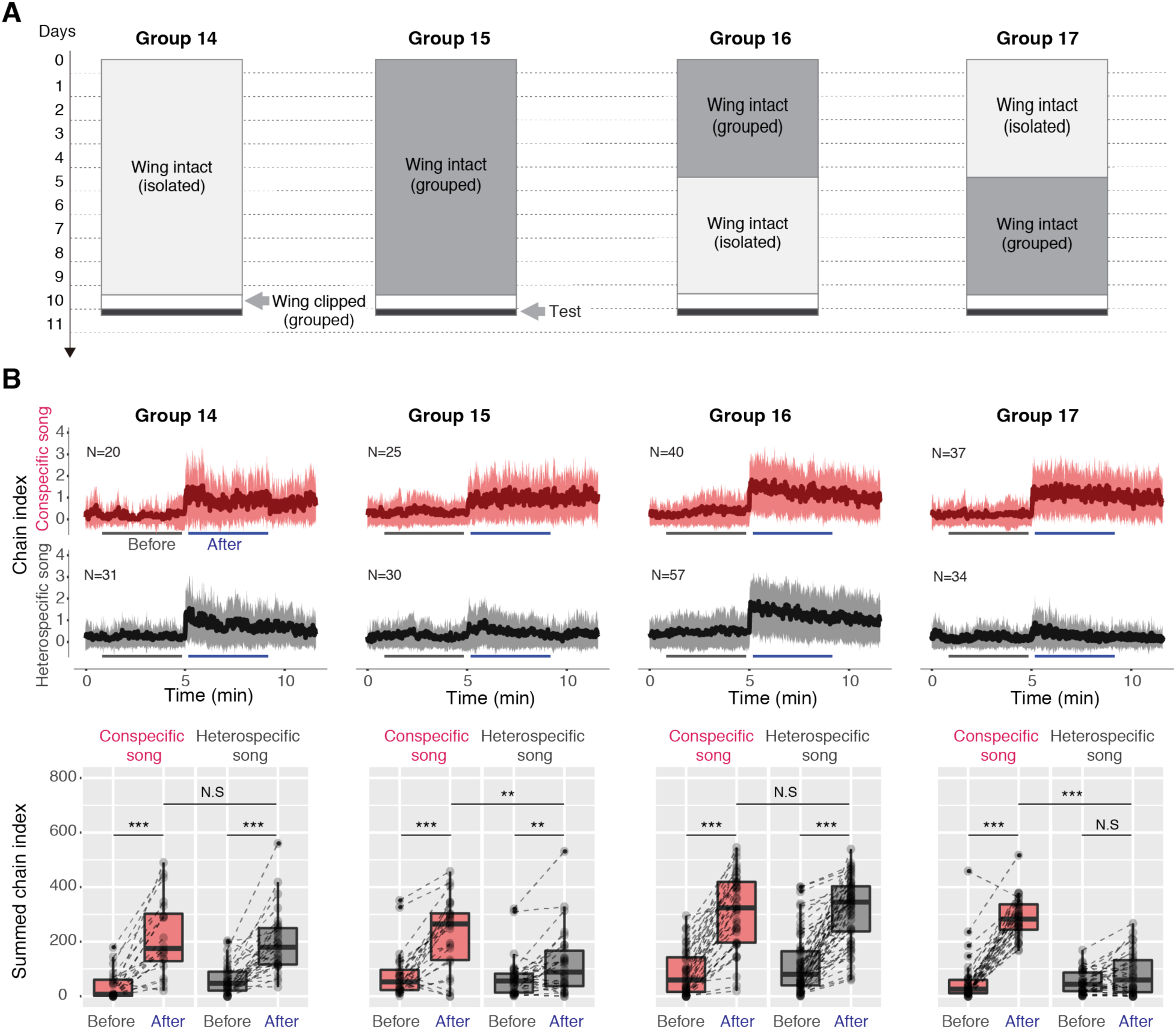
Lost song preference can be restored by auditory exposure. (A) Wing-intact flies were reared for 10 days after eclosion in either isolated (Group 14) or grouped conditions (Group 15) prior to the chaining tests. Flies in Group 16 were reared in grouped conditions for 5 days and then in isolated conditions for another 5 days. Flies in Group 17 were reared in the same conditions as Group 16, but in the reverse order. Refer to **Table S1** for the experimental conditions. (*B*) Time-courses of the chain index in response to playback of the conspecific song (red) and heterospecific song (grey). N.S., not significant, *P*>0.05; \**P*<0.05; \*\**P*<0.01; \*\*\**P*<0.001 Mann-Whitney U tests with the Benjamini–Hochberg correction and Wilcoxon signed-rank test with the Benjamini–Hochberg correction were used in multiple comparisons of mixed samples. N, number of behavioral chambers examined. Refer to **Table S2** for details of statistical tests and scores.

Flies isolated continuously for 10 days (Group 14) showed no song preference in the chaining tests (**Fig. 4*B***), while flies grouped for 10 days (Group15) exhibited a significant preference for the conspecific song (**Fig. 4*B***). Flies whose rearing conditions were switched from grouped to isolation on the 5th day (Group 16) showed no song preference (**Fig. 4*B***), which is consistent with the results shown in **Fig. 2** and strengthens the idea that maintaining song preference requires song input. On the other hand, when the order of the 2 rearing conditions was reversed (Group 17; 5 days isolation followed by 5 days of group conditions), the flies exhibited a song preference (**Fig. 4*B***). These results, together with the previous finding that flies have no song preference when isolated for 5 days after eclosion (**24**), suggest that male flies can reacquire song preference after the preference is lost.

We further tested if an extended isolation period eliminates preference recovery by extending the initial isolation period before rearing the flies in group conditions for 5 days (**Fig. 5*A***). We found that flies still exhibited a song preference even when the isolation period was extended to 10 days (Group 18, **Fig. 5*B***) or 15 days (Group 19, **Fig. 5*B***), suggesting a robust ability to reacquire song preference, even in aging flies, irrespective of the length of the isolation period.

**Fig. 5.**
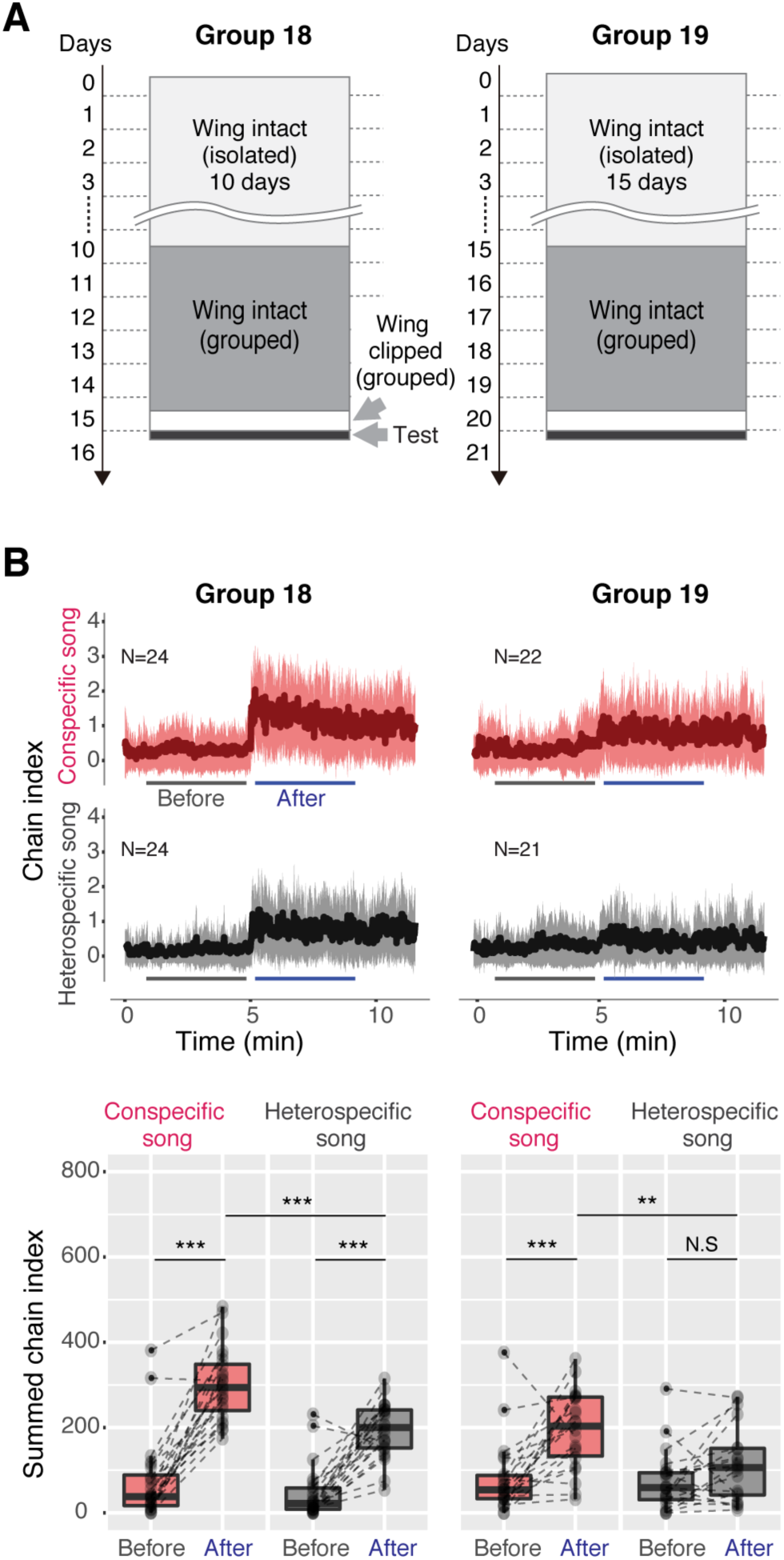
Lost song preference can be recovered over extended periods of time. (*A*) Flies in Group 18 were reared in isolated conditions for 10 days and then in grouped conditions for 5 days. Flies in Group 19 were reared in isolated conditions for 15 days and then in grouped conditions for 5 days. Refer to **Table S1** for the experimental conditions. (*B*) Time-courses of the chain index in response to playback of the conspecific song (red) and heterospecific song (grey). N.S., not significant, *P*>0.05; \**P*<0.05; \*\**P*<0.01; \*\*\**P*<0.001 Mann-Whitney U tests with the Benjamini–Hochberg correction and Wilcoxon signed-rank test with the Benjamini–Hochberg correction were used in multiple comparisons of mixed samples. N, number of behavioral chambers examined. Refer to **Table S2** for details of statistical tests and scores.

## Discussion

In auditory perceptual preference research, the mechanisms by which innate predisposition and experience interact remain elusive (**4, 37, 38**). While songbirds offer a powerful model for studying species-specific song discrimination, equivalent experiments are not feasible in humans and thus the focus has switched to phonetic discrimination between languages (**4, 39**). Progression in our understanding of auditory preference is hindered by the highly complex nature of auditory interactions in these traditional models. The establishment of a relatively simple model would thus greatly facilitate investigations of the underlying molecular and neural mechanisms. Here we propose the use of *Drosophila* as such a model organism, and provide a framework for the progression of song preference in *Drosophila* (**Fig. 6**).

**Fig. 6.**
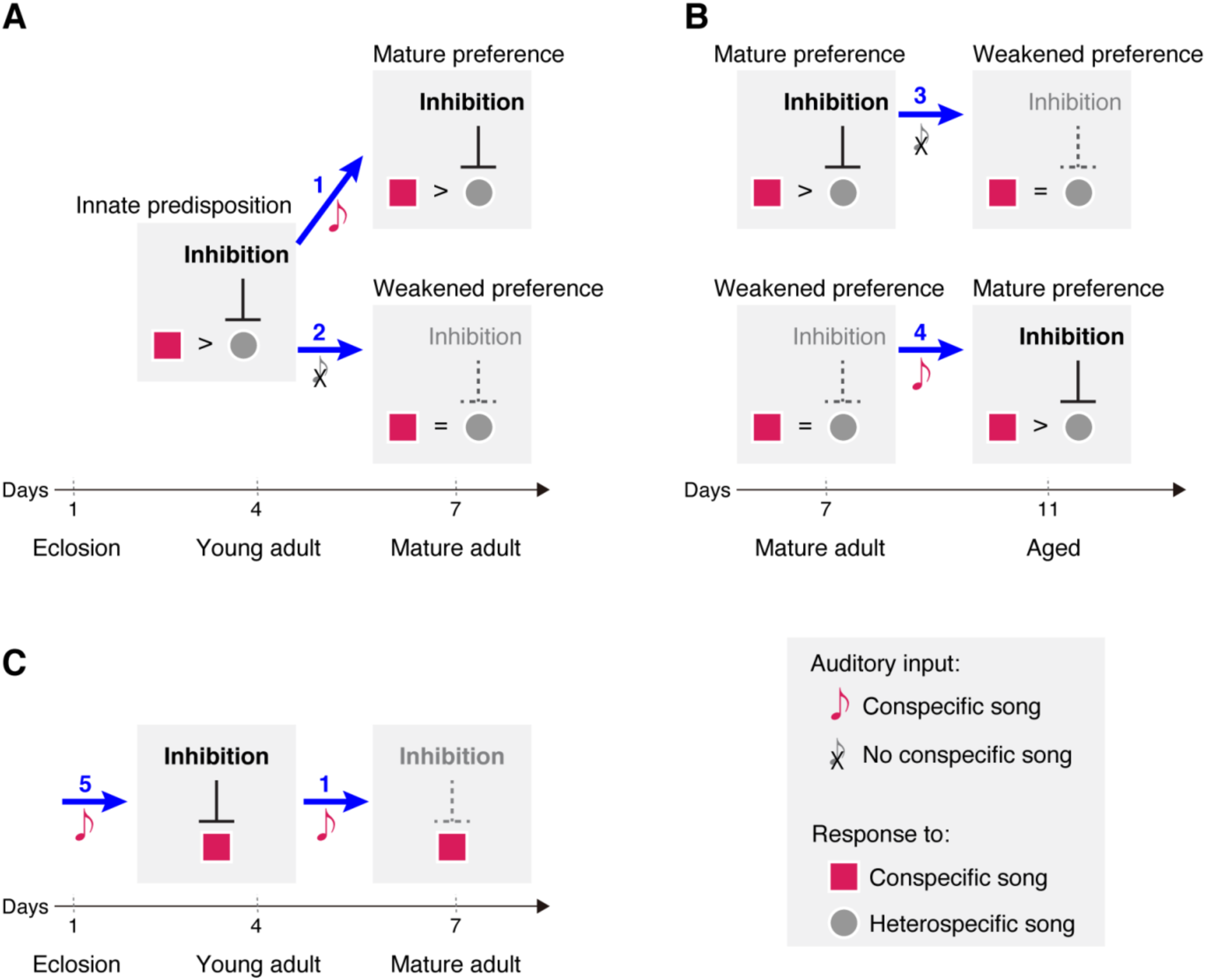
A framework of song preference progression in *Drosophila melanogaster*. Song exposure here specifically represents conspecific song exposure. Manipulation procedures are represented by numbers. In processes 1, 4, & 5, the flies are exposed to conspecific song. In processes 2 & 3, the flies are deprived of conspecific song experience. Only experience-dependent inhibition is discussed in this framework. (A) The song preference in young adult flies is independent of song experience, and should derive from innate predispositions. During the transition to a mature adult, maintenance of song preference depends on the conspecific song experience. (B) During aging, the song preference could be lost by conspecific song deprivation and restored by conspecific song exposure. (C) In young flies, the conspecific song response is inhibited by song experience but recovers when the flies mature. The experience-dependent plasticity of the conspecific song response thus differs from that of the heterospecific song response.

Both songbirds and humans possess innate predispositions for auditory learning. In songbirds, innate recognition of conspecific songs (**7, 8, 9**) and preference for learning conspecific songs (**6, 15**) are assumed to be independent of early auditory experience (**4, 15, 38, 40**), but rooted in genetic divergence (**6, 37, 41**). As an analogous model, fruit flies demonstrate an innate predisposition for song preference (**Fig. 1**). Further, we demonstrated that innate song preference cannot be sustained during periods of song deprivation or social isolation (**Figs. 2**, **4**, and **6*A***); auditory input reinforces innate song preference by preventing its decline. Specifically, song preference is maintained or restored only by exposure to conspecific song, but not to heterospecific song (**24**). These results favor a selective model of auditory learning in flies, reminiscent of similar findings in birds and humans (**4, 15**).

The ability of infants to acquire any language is accompanied by the ability to discriminate phonetic contrasts in all languages (**4**). The capacity to make such distinctions rapidly declines, however, if the individuals are not exposed to languages that express these distinctions (**1**). Here, our results show a similar phenomenon; young naive flies can discriminate conspecific songs from heterospecific songs (**Fig. 1**), while mature naive flies lose this ability (**24**; **Fig. S1**). Conspecific song experience can, however, help to maintain this discrimination ability during maturation (**Fig. 6*A***). On the other hand, in contrast to songbirds and humans, which have a relatively limited period of sensitivity to auditory acquisition, our study suggests that *D. melanogaster* has a robust ability to reacquire auditory preference during aging (**Fig. 6*B***). These similarities and dissimilarities in the plasticity of song preference between *Drosophila* and other models provide an attractive direction for research into the underlying neural mechanisms.

Although conspecific-selective neural activities are found in diverse regions in songbirds, including primary audition region field L, secondary auditory nuclei NCM & CMM, and even sensory-motor vocal nuclei (**8, 19, 38, 42, 43**), it remains unclear how the preference toward conspecific song emerges and develops at the neuronal and molecular levels (**37**). The rich genetic tools available for *D. melanogaster* and the short life cycle of flies would facilitate such investigation. In songbirds, song learning requires hearing experience, and experience-dependent maturation of synaptic inhibition is associated with selective auditory response (**43**) and maintenance of song production (**44**). A better understanding of how the maturation of inhibitory circuitry governs the development of song preference might help to elucidate the underlying mechanism. Indeed, maintenance of song preference in flies requires GABAergic input onto central courtship neurons (**24**), consistent with the above hypothesis. Thus, in this framework, we assume that experience-dependent inhibition represents suppression of the heterospecific song response (**Fig. 6**). Further studies are needed to explore the dynamic change in the inhibitory network.

Our present findings demonstrate the complex plasticity of auditory responses in *D. melanogaster* (**Fig. 6**). Song preference is governed by separate dynamic changes in response to conspecific and heterospecific songs, and the response to conspecific and heterospecific songs manifests both temporal and experience-dependent plasticity. First, the auditory response to conspecific songs in young naive flies (Group 1) was as strong as that in mature naive flies, while the auditory response to heterospecific songs in young naive flies was much weaker than that in mature naive flies (**Fig. S1**), suggesting that regulatory mechanisms underlying the responses to conspecific and heterospecific songs temporally differ. The innate predisposition might be manifested through selective inhibition of the response to heterospecific song. Second, an experience-dependent change in song preference is mainly mediated by the plastic response to heterospecific song (**Fig. 2**), probably involving GABAergic inhibition on the heterospecific song response (**24**). Deprivation of courtship-song hearing, either by wing-clipping or social isolation, weakened the conspecific song preference by releasing inhibition, thus increasing the response to the heterospecific song (**24**; **Fig. 6*A***). The weakened song preference could be strengthened by re-exposure to conspecific songs during a relatively long sensitive period (**Figs. 4, 5**, and **6*B***). In contrast, conspecific song input maintained the innate predisposition and manifested it as a mature preference (**Figs. 1**, **4**, and **6*A*; 24**). The maintained song preference in the mature adult could be lost again, however, if deprived of conspecific song input (**Figs. 2**, **4**, and **6*B***). Our model suggests the existence of a complex interaction between genetic factors and experience in shaping the song preference in *Drosophila*.

The experience-dependent plasticity of the conspecific song response differs from that of the heterospecific song response. Although auditory experience of conspecific songs suppressed the response to heterospecific songs in all observed life stages (**24**; **Fig. 6 *A*** and ***B***), the response to the conspecific song was also reduced in young, but not mature, flies (**24**; **Figs. 3** and **6*C***). Importantly, the experience-dependent reduction in the response to conspecific song in the young adult did not change the preference to the conspecific song over the heterospecific song (**Fig. 1*C***). Some unknown mechanisms that underlie the innate song preference may regulate the responses to both conspecific and heterospecific songs, and such regulation could be vulnerable to song input in young adults. Further investigation is needed to determine whether the hypothesized inhibition that underlies innate song preference is shared with experience-dependent inhibition. Importantly, these mechanisms together constitute and secure song preference.

A study in which song-induced locomotion was measured in *D. melanogaster* showed that when tested singly, wing-intact flies reared in an isolated condition, which are considered to be naive flies, respond only to conspecific songs, and not to heterospecific songs when aged for 5-7 days (**45**). When tested in a group via chaining assays, however, naive flies of the same age that were also reared in an isolated and wing-intact condition show identical responses to conspecific (35 ms IPI) and heterospecific songs (75 ms IPI) (**23, 24**). This discrepancy possibly derives from the different experimental conditions; in the chaining assay, 6 flies were grouped in a chamber and they interacted with each other during testing of their song preferences. In such group-testing environments, acoustic information would be integrated with information of visual and tactile stimuli from other flies, which eventually reduces response thresholds and increases the sensitivity of the auditory response of flies. Further physiologic studies are required to demonstrate the response selection on a neuronal level.

Song preference is a fundamental mechanism for reproductive isolation for many animals, including *Drosophila* (**24, 46, 47, 48**). Both an innate bias toward conspecific songs and experience-dependent reinforcement could contribute to avoiding interspecies mating. Our results explain the interaction of experience and innate song preference on a behavioral level. Taken together, our study provides a complementary model to songbirds and humans for future investigation into the neural mechanisms underlying the interactions between auditory experience and innate sound predisposition.

## Materials and Methods

### Animals

The wild-type strain Canton-S of *D. melanogaster* was raised on standard yeast-based media at 25°C in 40% to 60% relative humidity on a 12 h light/ 12 h dark cycle. Adult male flies were used for all the experiments.

### Fly preparations

Virgin males were collected within 8 h of eclosion. Days after eclosion are used to indicate the age of flies. Since male flies produce courtship songs by unilateral wing vibrations (**31**), we clipped their wings to prevent them from emitting the song. The majority of the wing length was ablated but it did not affect the motor control of the wing root. For the chaining assay, the males had their wings clipped on the day of eclosion or one to four days before the chaining test. Flies were kept in groups of 6 to 8 males or singly for the chaining assay and 10 to 12 for the homosexual courtship recordings. Each condition is described in **Table S1** and illustrated accordingly in each figure.

### Recording of homosexual courtship behavior

Virgin males were kept in groups for 2, 3, or 4 days after eclosion with their wings intact. We used a round chamber with a diameter of of 90 mm, which has the same sloped-wall design of the flybowl chamber described previously, for the behavioral recordings (**49**). 9 males were gently introduced to the chamber and their behaviors were recorded at 25°C in 50% to 70% relative humidity. The flies’ contours were outlined by a backlit LED light box (Maxon Tracer LED LD-A4, Holbein Art Materials Inc, Tokyo, Japan) and captured by a CMOS camera (DFK33UP1300, The Imaging Source Asia Co., Ltd., Taiwan) for 30 min at 30 fps. The recording was performed between 1 and 3 h after light onset of the light / dark cycle.

### Sound stimuli used in the training and chaining assay

We define “training” as the process that exposes flies to external artificial sounds (**24**). In *D. melanogaster*, the natural pulse song has a species-specific IPI distributed between 29 and 46 ms with a mean of 35-36 ms (**50**). Artificial pulse songs with 35 ms and 75 ms IPIs were thus used as conspecific and heterospecific songs, respectively, for training and chaining assay (**24**). In the training, we used the conspecific song, white noise, and a pure tone of 167 Hz (**Table S1**), with a mean baseline-to-peak amplitude of the particle velocity between 6.6 and 8.6 mm/s as described previously (**24**). Mean baseline-to-peak amplitude of the particle velocity was 9.2 mm/s for the sound stimuli used in the chaining assay (**51**).

### Exposure to artificial sounds

Flies in Groups 9 and 10 were trained with white noise and pure tones (167 Hz), respectively. Flies in Groups 11, 12 & 13 were trained with conspecific song. Each condition is described in **Table S1** and illustrated in **Fig. 3*A***. The training method is the same as the protocol described previously (**33**). After the training session, the trained flies were kept in groups of 6 to 8 males until the chaining assay.

### Chaining assay to quantify the auditory response

We used a chaining assay to examine the auditory response of male flies (**23, 24, 52**). The test was performed between 0 and 3 h after light onset of the light / dark cycle. Six flies were loaded into one lane of an acoustic behavior chamber (**32**) and placed in front of a loudspeaker at a distance of around 11 cm. A sound stimulus was delivered from a loudspeaker with an amplifier (Lepai LP-2020A + NFJ Edition, Bukang Electrics, Jieyang, China). The flies’ contours were outlined by a backlit LED light box (ComicMaster Tracer, Too Marker Products, Tokyo, Japan), and captured by a monochrome camera (Himawari GE60, Library, Tokyo, Japan) with a zoom lens (Lametar 2.8/25 mm, Jenoptik GmbH, Jena, Germany). Flies were not exposed to sound for the first 5 min, and were then exposed to a sound stimulus that lasted for 6.5 min. The recorded video was then down-sampled to 1 Hz and analyzed off-line using ChaIN version 3 (**51**), which measured only the number of follower flies in chains as the chain index. To compare the chain index before and after the song onset, summed chain index was calculated by taking the sum of chain index 4-min before (51-290 s) and after (311-550 s) the song onset (indicated by the horizontal lines). Increase in the chain index following song onset was defined as the difference between the summed chain index from 311-550 s and the summed chain index from 51-290 s.

### Statistical analyses

Statistical analysis was performed using R (R Core Team, version 3.5.3) (**53**). We identified that some of our data were not normally distributed (Shapiro-Wilk test for normality), and thus non-parametric statistical analyses were applied. For multiple comparison of mixed samples, Mann-Whitney U tests with Benjamini and Hochberg (BH) correction were used to compare the summed chain index of two independent samples, and Wilcoxon signed-rank test with BH correction was used to compare the chaining behavior before and after the sound onset. To compare more than two groups of independent samples, a Kruskal–Wallis test followed by the Mann-Whitney U test with BH correction was used. Detailed statistical results are shown in **Table S2**. Boxplots were drawn using the ggplot2 package (version 3.2.1) in R. Boxplots display the median of each group along with the 25th and 75th percentiles. Whiskers denote 1.5x the inter-quartile range.

## Supporting information

Table S1. Summary of all experimental conditions for each group tested.

Table S2. Summary of all statistical results for each comparison.

Movie S1. Spontaneous homosexual courtship of 2-day-old male flies.

Movie S2. Spontaneous homosexual courtship of 3-day-old male flies.

Movie S3. Spontaneous homosexual courtship of 4-day-old male flies

## Acknowledgments

We thank N. Aisaka, M. Kuno, and Y. Maki for fly maintenance, and M. P. Su, R. Tanaka, and K. Wada for discussion. This work was supported by MEXT KAKENHI Grants-in-Aid for Scientific Research (B) (JP20H03355 to AK and JP18H02488 to YI), Grant-in Aid for Scientific research (C) (15K07147 and 18K06332 to HI), Scientific Research on Innovative Areas “Evolinguistics” (Grant JP18H05069 and JP20H04997 to AK), “Systems science of bio-navigation (Grant JP19H04933 to AK), and “Constrained and directional evolution” (Grant JP18H04819 to YI), Challenging Research (Exploratory) (JP17K19450 to AK and JP17K19425 to YI), the Naito foundation to AK, Inamori Foundation Research Grant, Japan to HI, and Grant-in-Aid for JSPS research Fellow (18J15228 to XL).

## Supplementary Appendix

**Fig. S1.**
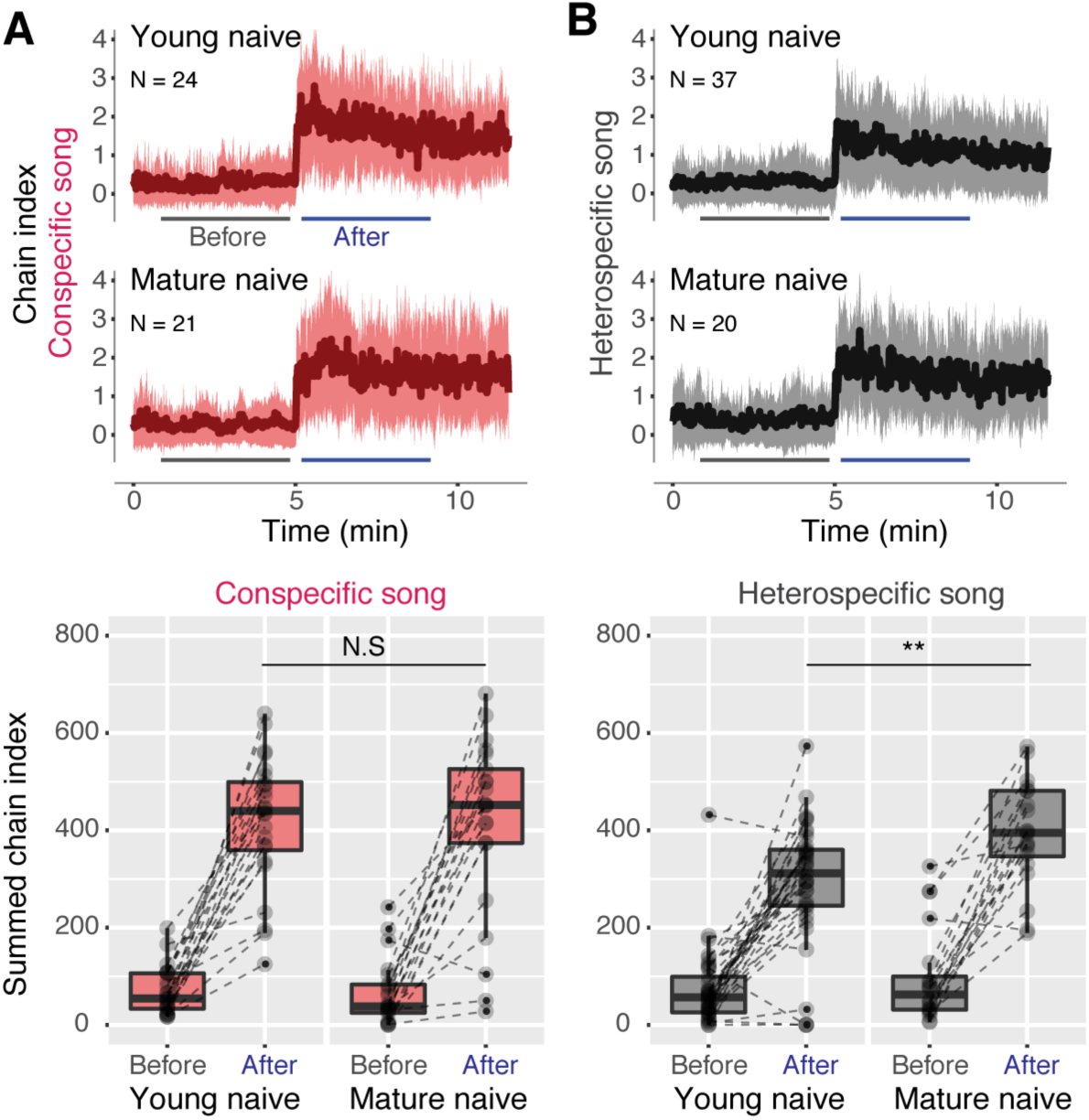
Temporal differentiation of auditory response to songs. (*A*) Auditory response to conspecific songs for 4-day-old males (Young naive; Group 1 in the present study) and 7-day-old males (Mature naive; data from Li et al. [**24]**, group-reared). (*B*) Auditory response to heterospecific songs for 4-day-old males (Young naive; Group 1 in the present study) and 7-day-old males (Mature naive; data from Li et al. [**24**], group-reared). N.S., not significant, *P*>0.05; \**P*<0.05; \*\**P*<0.01; \*\*\**P*<0.001 Mann-Whitney U tests with the Benjamini–Hochberg correction and Wilcoxon signed-rank test with the Benjamini–Hochberg correction were used in multiple comparisons of mixed samples. N, number of behavioral chambers examined. Refer to **Table S2** for details of statistical tests and scores.

**Fig. S2.**
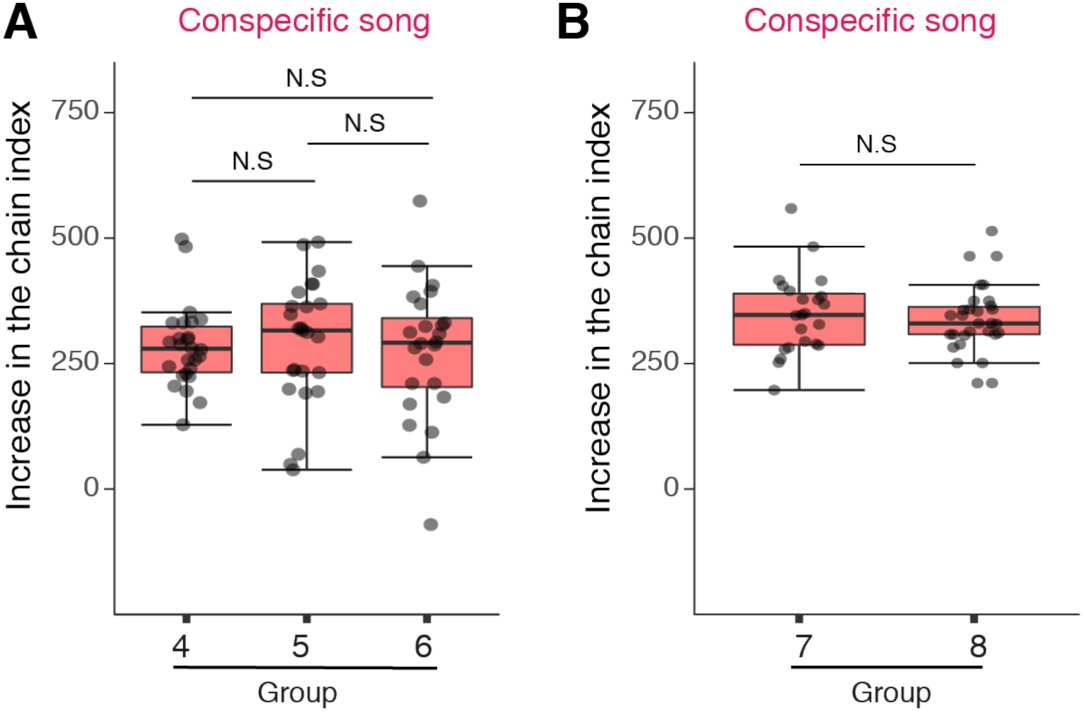
Increase of chaining activities during conspecific song stimulation for Groups 4, 5 & 6 (*A*), and for Groups 7 & 8 (*B*). Increase in the chain index following song onset = (summed chain index in 311-550 sec) – (summed chain index in 51-290 sec). N.S., not significant, *P*>0.05; \**P*<0.05; \*\**P*<0.01; \*\*\**P*<0.001 Kruskal–Wallis tests followed by the Mann-Whitney U test with the Benjamini– Hochberg correction was used in multiple comparisons of independent samples; Mann-Whitney U tests were used for 2-sample comparisons. N, number of behavioral chambers examined. Refer to **Table S2** for details of statistical tests and scores.

**Fig. S3.**
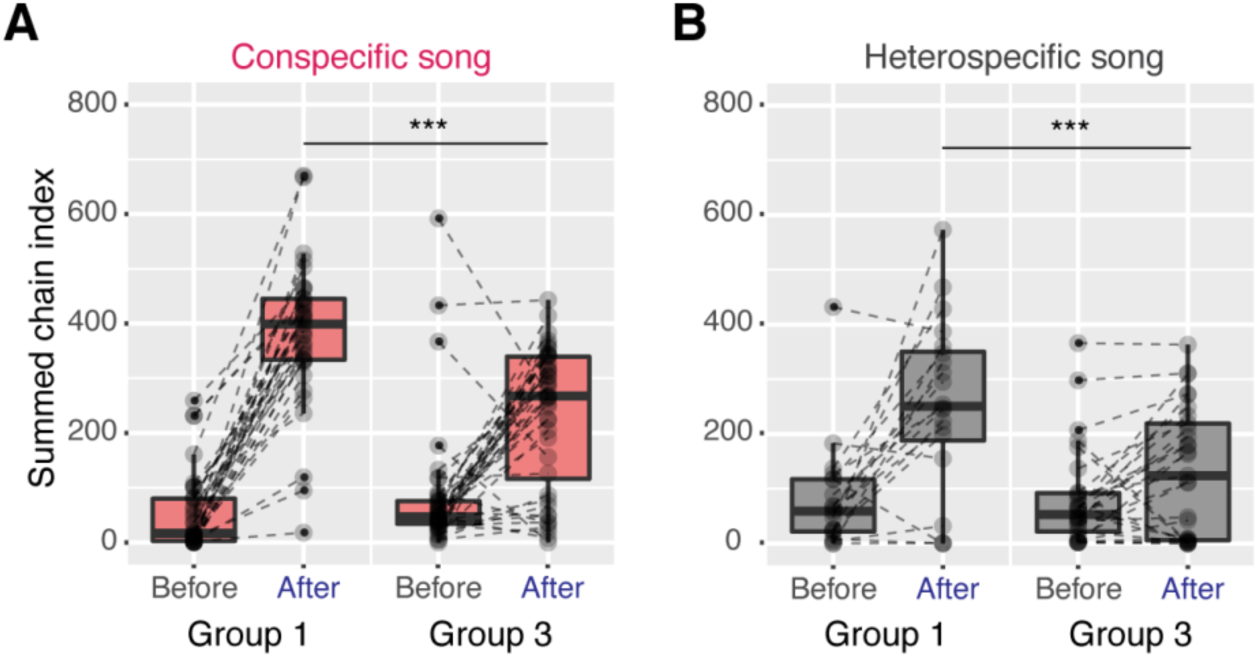
Young intact males show reduced auditory responses compared to wing-clipped males. (*A*) Auditory responses to conspecific songs for wing-clipped young males (Group 1) and intact young males (Group 3). (*B*) Auditory responses to heterospecific songs for wing-clipped young males (Group 1) and intact young males (Group 3). N.S., not significant, *P*>0.05; \**P*<0.05; \*\**P*<0.01; \*\*\**P*<0.001 Mann-Whitney U tests with the Benjamini–Hochberg correction and Wilcoxon signed-rank test with the Benjamini–Hochberg correction were used in multiple comparisons of mixed samples. N, number of behavioral chambers examined. Refer to **Table S2** for details of statistical tests and scores.

## SI Appendix Tables

**Table S1**. Summary of all experimental conditions for each group tested.

**Table S2**. Summary of all statistical results for each comparison.

## SI Appendix Movies

**Movie S1-S3**. Typical spontaneous homosexual courtship of 2-4 days old male flies, respectively. In a 30-min observation window, spontaneous chasing behavior and unilateral wing extension between males were observed.

